# Structural basis for binding mechanism of human serum albumin complexed with cyclic peptide dalbavancin

**DOI:** 10.1101/2020.09.09.287375

**Authors:** Sho Ito, Akinobu Senoo, Satoru Nagatoishi, Masahito Ohue, Masaki Yamamoto, Kouhei Tsumoto, Naoki Wakui

## Abstract

Cyclic peptides, with unique structural features, have emerged as new candidates for drug discovery; their association with human serum albumin (HSA; long blood half-life), is crucial to improve drug delivery and avoid renal clearance. Here, we present the crystal structure of HSA complexed with dalbavancin, a clinically used cyclic peptide. SAXS and ITC experiments showed that the HSA-dalbavancin complex exists in a monomeric state; dalbavancin is only bound to the subdomain IA of HSA in solution. Structural analysis and MD simulation revealed that the swing of Phe70 and movement of the helix near dalbavancin were necessary for binding. The flip of Leu251 promoted the formation of the binding pocket with an induced-fit mechanism; moreover, the movement of the loop region including Glu60 increased the number of non-covalent interactions with HSA. These findings may support the development of new cyclic peptides for clinical use, particularly the elucidation of their binding mechanism to HSA.

## INTRODUCTION

Cyclic peptides are attracting increasing attention not only in the pharmaceutical industry but also in academic research. The establishment of high-speed screening systems rendered cyclic peptides particularly attractive for drug discovery. High-speed screening systems developed to date include phage display^1,2^ and SICLOPPS^3,4^ as *de novo* discovery approaches based on *in vivo* production. Moreover, Passioura et al. developed another method called RaPID^5^ based on mRNA display. This system enables screening of very large numbers (up to 10^14^) of unique sequences; for the sake of comparison, phage display and other systems based on in *vivo* production can screen ‘only’ 10^8^-10^9^. RaPID also allows screening of cyclic peptides containing D-amino acids, β-amino acids, main chain modifications, and unnatural amino acids.^6,7^ Of note, the screening of cyclic peptides containing natural as well as non-natural amino acids from a large number of combinations enabled the identification of peptides showing a high binding capacity to various target proteins.^8-11^ While conventional linear peptides are highly unstable against gastric acid pH, cyclization allows the overcoming of this problem suggesting that cyclic peptides may be orally administered for therapeutic purposes. Furthermore, although conventional peptide pharmaceuticals are challenged by proteolytic degradation, the main chain structure modifications and unnatural amino acids contained in cyclic peptides render them less susceptible to degradation. Currently, more than 40 cyclic peptides have been approved for human use and 20 others are in the development stage.^12^

Nevertheless, cyclic peptide drug discovery is also facing some challenges, particularly concerning absorption, distribution, metabolism, excretion, and toxicity (ADMET). For example, renal excretion is an issue in the development of cyclic peptides as therapeutics. In general, exogenous peptides are rapidly filtered by the kidneys. The glomeruli have a porous structure and efficiently filter plasma molecules with a molecular weight lower than ~5-10 kDa.^13^ Additionally, as cyclic peptides are also less likely to be reabsorbed from the renal tubules, they rapidly undergo renal clearance. Binding to human serum albumin (HSA) is, therefore, important to avoid renal clearance, because this 66 kDa protein is not affected by glomerular filtration and has a very long blood half-life (~20 days). Therefore, methods aimed at increasing the binding affinity to albumin have been developed and used for conventional peptide drugs. For instance, the conjugation with albumin using a chemical linker was used for exendin-4-albumin conjugate (CJC-1143-PC) and GLP-1. Other studies successfully avoided peptide drugs’ renal clearance by conjugation to a molecule that is known as an albumin-binder.^14-18^

Currently, no structure of a cyclic peptide in complex with HSA is available in Protein Data Bank (PDB). Structures of HSA registered in PDB include complexes with small molecules.^19-21^ These structures revealed major binding sites, such as Sudlow’s sites for most drug molecules, and fatty acid binding sites for fatty acids. However, the mechanism by which cyclic peptides interact with HSA remains poorly understood. Therefore, structural information on the binding site of cyclic peptides in albumin is required to successfully avoid their renal excretion. Dalbavancin, along with other similar drugs such as telavancin, oritavancin, teicoplanin, and vancomycin, is a cyclic lipoglycopeptide containing a common heptapeptidic core with five residues. Their unique structural features, such as a hydrophilic cyclic moiety and a hydrophobic hydrocarbon chain, raise the question of whether dalbavancin binds to Sudlow’s sites, fatty acid binding sites, or an entirely new binding site. Having a high plasma protein-binding ability (~93-98%) and a long half-life (~6-11 days), dalbavancin is a good candidate to obtain a structure of HSA complexed with a cyclic peptide.^22-24^ Dalbavancin is used for the treatment of complicated skin infections, bloodstream infections, endocarditis, bone and joint infections, and/or meningitis caused by methicillin-resistant *Staphylococcus aureus* or other Gram-positive infections. Importantly, the structural insights provided by HSA-cyclic peptide complex may elucidate the problem of renal clearance of cyclic peptides already in the initial stages of high-speed screening. Here, we report the first three-dimensional structure of HSA in complex with dalbavancin.

## RESULTS AND DISCUSSION

### Crystal structure of HSA with/without dalbavancin

We determined the crystal structure of HSA with dalbavancin at a resolution of 2.8 Å and that of the HSA apo form at a resolution of 2.6 Å (Table 1). The apo form structure of HSA is similar to the previously reported ligand-free structure (PDB ID: 4K2C).^25^ Electron density from phosphate ion was clearly observed at Drug site 1, also known as Sudlow’s site.^26^ In the dalbavancin-bound structure, there were three HSA molecules in the asymmetric unit. Each molecule had a similar structure (RMSD using all atoms of each chain, chain A vs chain B: 0.825 Å, chain A vs chain C: 0.623 Å, and chain B vs chain C: 0.933 Å). Electron density from Jeffamine was observed at Drug site 1. The electron density from another Jeffamine was observed at the interface between subdomains IIA and IIB in a binding cleft that overlapped with the fatty acid-binding site known as FA6.^20,21^ The binding site of Jeffamine overlapped with that of ligands shown in other HSA structures.^26^ Clear electron density from dalbavancin was observed and located near the subdomains IA and IIIB (Figure 1a). Dalbavancin molecules (chain F, chain G, and chain H) were bound to the subdomain IA of each HSA in the asymmetric unit, showing binding interactions within each molecule. Moreover, each cyclic moiety of dalbavancin in the subdomain IIIB (chain I and J) bound to neighboring HSA (Figure S1b, Figure S1c, Figure S1d, and Figure S1e). Surprisingly, the cyclic moiety of dalbavancin in the subdomain IIIB displayed quite a different conformation from that of dalbavancin in the subdomain IA (Figure 1b). This indicates that the cyclic moiety of dalbavancin can assume a rather flexible structure.

**Table 1.** Data collection and refinement statistics

| PDB ID | Dalbavancin complex<br>6M5E | Apo<br>6M5D |
| --- | --- | --- |
| **Data collection** |  |  |
| Space group | C 2 | P 4 2<sub>1</sub> 2 |
| Cell dimensions |  |  |
| a, b, c (Å) | 197.05, 125.68, 125.74 | 182.70, 182.70, 79.81 |
| $\alpha, \beta, \gamma$ (°) | 90, 90.32, 90 | 90, 90, 90 |
| Resolution (Å)* | 42.82-2.80 (2.90-2.80) | 42.78-2.60 (2.70-2.60) |
| R<sub>meas</sub>* | 0.479 (5.311) | 0.078 (1.317) |
| R<sub>pim</sub>* | 0.101 (1.133) | 0.027 (0.451) |
| $\langle I/\sigma(I) \rangle$* | 9.28 (0.91) | 16.92 (1.59) |
| CC<sub>1/2</sub>* | 0.993 (0.456) | 0.999 (0.645) |
| Completeness(%)* | 99.9 (99.9) | 91.78 (94.01) |
| Redundancy* | 22.0 (21.7) | 7.9 (8.0) |
| **Refinement** |  |  |
| Resolution (Å) | 42.82-2.80 | 42.78-2.60 |
| No. unique reflections* | 75476 (7487) | 38650 (3891) |
| R<sub>work</sub> / R<sub>free</sub> | 0.1799 / 0.2276 | 0.2308 / 0.2585 |
| No. atoms |  |  |
| Protein | 13840 | 4470 |
| Ligand | 640 | — |
| water | 3 | 3 |
| Averaged B-factors(Å<sup>2</sup>) |  |  |
| Protein | 63.29 | 115.74 |
| Ligand | 85.35 | — |
| Water | 57.04 | 66.92 |
| R.m.s. deviations from ideal |  |  |
| Bond lengths (Å) | 0.18 | 0.01 |
| Bond angles (°) | 1.49 | 1.27 |
| Ramachandran plot |  |  |
| Favored (%) | 95.91 | 90.50 |
| Allowed (%) | 3.34 | 6.63 |
| Outlier (%) | 0.75 | 2.87 |

**Figure 1.**
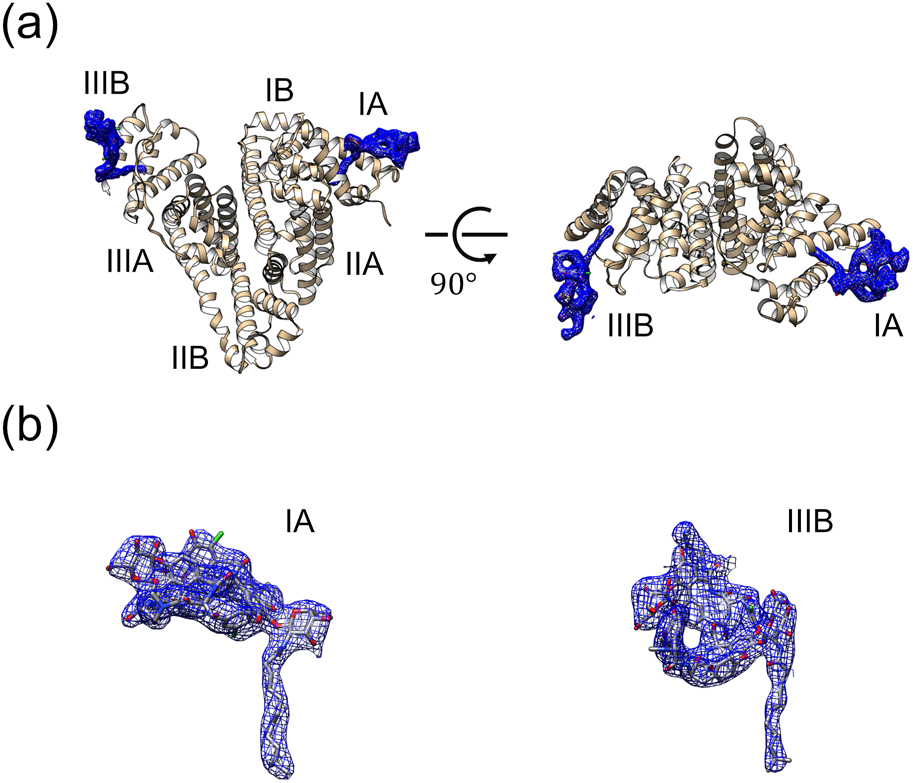
Crystal structure of the HSA-dalbavancin complex and dalbavancins. (a) Overall crystal structure (tan) of the HSA-dalbavancin complex (PDB ID: 6M5E). The name of each subdomain is shown in the figure. 2mFo-DFc map of bound dalbavancins contoured at 1.0 σ. Two dalbavancins were bound per HSA molecule in the crystal structure. One of the dalbavancins (left side of each figure) interacted with the neighboring HSA via its cyclic moiety. (b) A close-up view of dalbavancins in each subdomain.

### Dalbavancin binds only to the subdomain IA in solution

The interaction of the cyclic moiety of dalbavancin in subdomain IIIB with the neighboring HSA (chain B) observed in the crystal structure (Figure S1) raises the question of how the HSA-dalbavancin complex exists in solution. In fact, if the interaction observed in the crystal structure was similar in solution, the HSA-dalbavancin complex could not maintain monodispersity. Therefore, we determined the state of assembly of HSA (5 mg mL^−1^) in solution with 400 μM dalbavancin (Table S1 and Figure S3). The Rg values of the crystal structure (using chain C with chain H and chain I for calculation with CRYSOL), and of the solution structure were 27.9 Å and 28.2 Å, respectively (Table S1), indicating that the HSA-dalbavancin complex exists in a monomeric state in solution (Figure 2a). The resultant structure also implies the following states of dalbavancin in subdomain IIIB: 1. dalbavancin shows different conformations from those seen in the crystal structure and the cyclic moiety cannot form strong enough interactions with neighboring HSA molecules, or 2. dalbavancin cannot bind to the subdomain IIIB. Nonetheless, it is still unclear whether dalbavancin binds to subdomain IIIB in solution or not. Next, we determined the stoichiometry between HSA and dalbavancin using isothermal titration calorimetry (ITC). ITC showed that the molar ratio of dalbavancin against one HSA molecule was approximately one (n = 0.84), implying that the stoichiometry between HSA and dalbavancin in solution is likely 1 : 1 (Figure 2b, Figure S6, and Table S2). The binding of dalbavancin to HSA was enthalpy-driven (Δ*H* = −8.9 kcal mol^−1^, −*T*Δ*S* = 1.9 kcal mol^−1^), which seems to reflect the multiple hydrogen bonds between the cyclic moiety of dalbavancin and HSA as shown in Figure 3b. Indeed, the binding of dalbavancin to HSA was quite strong; an exothermic reaction upon binding of dalbavancin was observed in the presence of lauric acid, a competing fatty acid, suggesting that dalbavancin may bind to HSA in an environment such as blood circulation which exists various fatty acids (Figure S4). Then, to determine which of the subdomains, IA or IIIB, is the binding site of dalbavancin in solution, further ITC experiments were performed using recombinant human serum albumin domain I-II or domain III. When either domain I-II or domain III was titrated with dalbavancin, an enthalpy-driven reaction was observed only in the condition using domain I-II (Figure 2c, Figure 2d, Figure S5, and Figure S6). The molar ratio and thermodynamic parameters of the binding of dalbavancin to domain I-II were comparable to those for the binding of dalbavancin to full-length HSA (Table S2). These data indicate that the binding site of dalbavancin to HSA in solution is subdomain IA.

**Figure 2.**
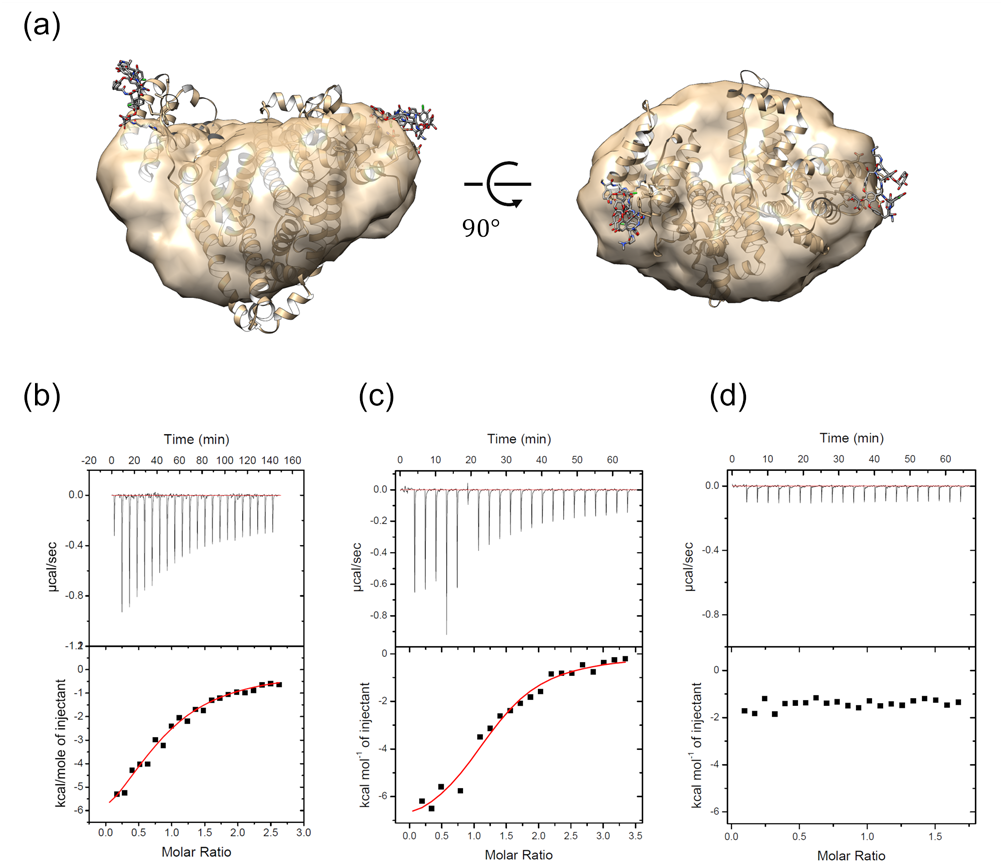
Binding characterization of dalbavancin and HSA in solution. (a) HSA-dalbavancin complex (tan). Crystal structure of the HSA-dalbavancin complex (represented by ribbon models) is superimposed to the SAXS models (represented by surface models). SAXS analysis revealed that HSA exists as a monomer in solution (see also Table S1). (b-d) Thermodynamic characterization of dalbavancin interaction with full-length HSA, domain I-II, and domain III by ITC. The top and bottom panels show the titration kinetics and the binding isotherm, respectively. The fitting curve with the program ORIGIN7 is indicated with a red trace (bottom panel). b) Dalbavancin (400 μM) was titrated into the cell that contained full-length HSA (25 μM). c) Dalbavancin (500 μM) was titrated into the cell that contained domain I-II (25 μM). d) Dalbavancin (500 μM) was titrated into the cell that contained domain III (50 μM).

**Figure 3.**
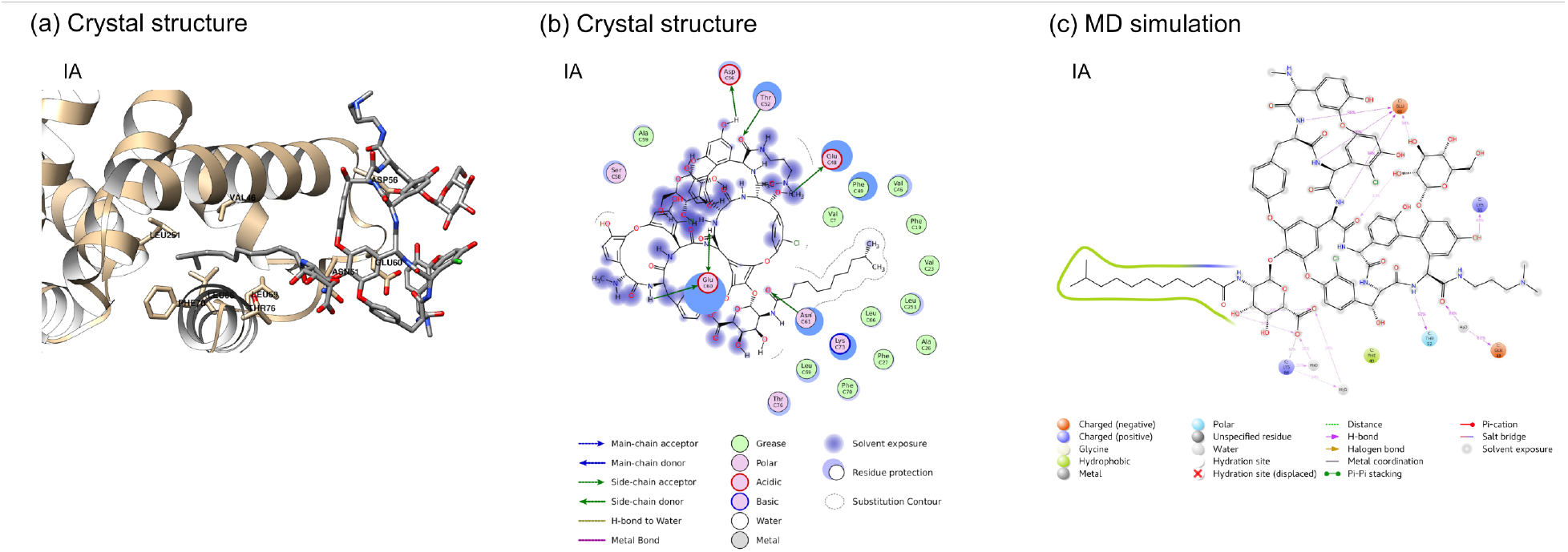
Interaction between HSA and dalbavancin. (a) The dalbavancin-binding pocket in subdomain IA (chain C) with dalbavancin (PDB ID: 6M5E). (b) The two-dimensional schematic diagram of the binding pocket. H-bonds (main chain, side chain, water-bridged) and amino acid types are represented by their symbols as displayed in the key. (c) The two-dimensional schematic diagram of interactions between dalbavancin and HSA from MD simulations. Interactions and amino acid types are represented by their symbols as displayed in the key.

### Dalbavancin in subdomain IA

The hydrocarbon chain of dalbavancin in subdomain IA of HSA was inserted into a hydrophobic pocket that comprises all nonpolar amino acids (Figure 3a and 3b), including Val7, Phe19, Val23, Phe27, Phe49, Leu66, Leu69, Phe70, and Leu251. Consequently, the highly hydrophobic environment of this pocket allows access to the hydrocarbon chain of dalbavancin.

The hydrophobic pocket in the subdomain IA has two remarkable structural features: 1. the swing of Phe70 and 2. the movement of helix 4 of subdomain IIA and flip of Leu251. Compared to the apo form, the helix 4 of the subdomain IA was slid of about 3 Å to accommodate the hydrocarbon chain of dalbavancin (Figure 4a). Interestingly, Phe70 was swung so that its side chain did not clash with the tail of the hydrocarbon chain and the hydrophobic binding pocket could be formed (Figure 4b). The representative structures obtained from the MD trajectory of dalbavancin-removed HSA showed Phe70 in a similar orientation as in the apo structure (Figure 4c). The hydrophobic binding pocket formed for the hydrocarbon chain of dalbavancin became smaller due to the swing of Phe70. Comparison with structures of HSA bound to different kinds of fatty acids revealed that HSA formed an actual binding pocket specific for the hydrocarbon chain of dalbavancin (Figure 5a). In the previously reported structures of lauric acid-bound and myristic acid-bound HSA^27^, both fatty acids interacted with the boundary region between subdomains IA and IIA as well as with the subdomain IA (Figure 5b and 5c). The environments surrounding the hydrocarbon chain of dalbavancin and lauric or myristic acid in the subdomain IA appear similar. Five hydrophobic residues (Val46, Phe49, Leu66, Leu69, and Phe70) are similarly oriented around these hydrocarbon chains. Moreover, the other lauric or myristic acid molecule, which was bound to the boundary region, contributed to form the hydrophobic environment for the lauric or myristic acid molecule in the subdomain IA. However, in order to fill the space occupied by the fatty acid in the boundary region as seen in lauric and myristic acid-bound HSA, helix 4 of the subdomain IIA (Leu250-Gln268) of HSA-dalbavancin complex moved approximately 3Å towards subdomain IA. Additionally, Leu251 from helix 4 of IIA flipped toward the hydrocarbon chain of dalbavancin and formed a hydrophobic environment (Figure 5a). Thus, the swing of Phe70, the movement of helix 4 from IIA, and the flip of Leu251 are key factors to form the binding pocket in the subdomain IA.

**Figure 4.**
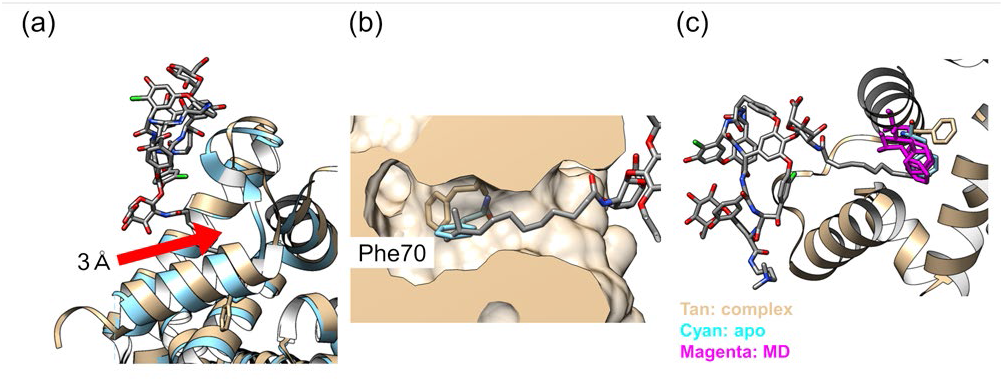
Structural differences between the complex (PDB ID: 6M5E) and the apo form (PDB ID: 6M5D). (a) Superimposed structures of the complex (tan) and the apo form (cyan) in the subdomain IA. (b) A close-up view of the binding pocket in subdomains IA. Phe70 of the complex (tan) swung not to overlap with the alkyl tail of the dalbavancin. However, the structure of Phe70 in the apo form (cyan) overlapped. (c) The structural alignment of the complex (tan), the apo form (cyan), and the representative structures of dalbavancin-removed HSA from each MD run (magenta). Phe70 from apo form and MD runs showed similar orientations.

**Figure 5.**
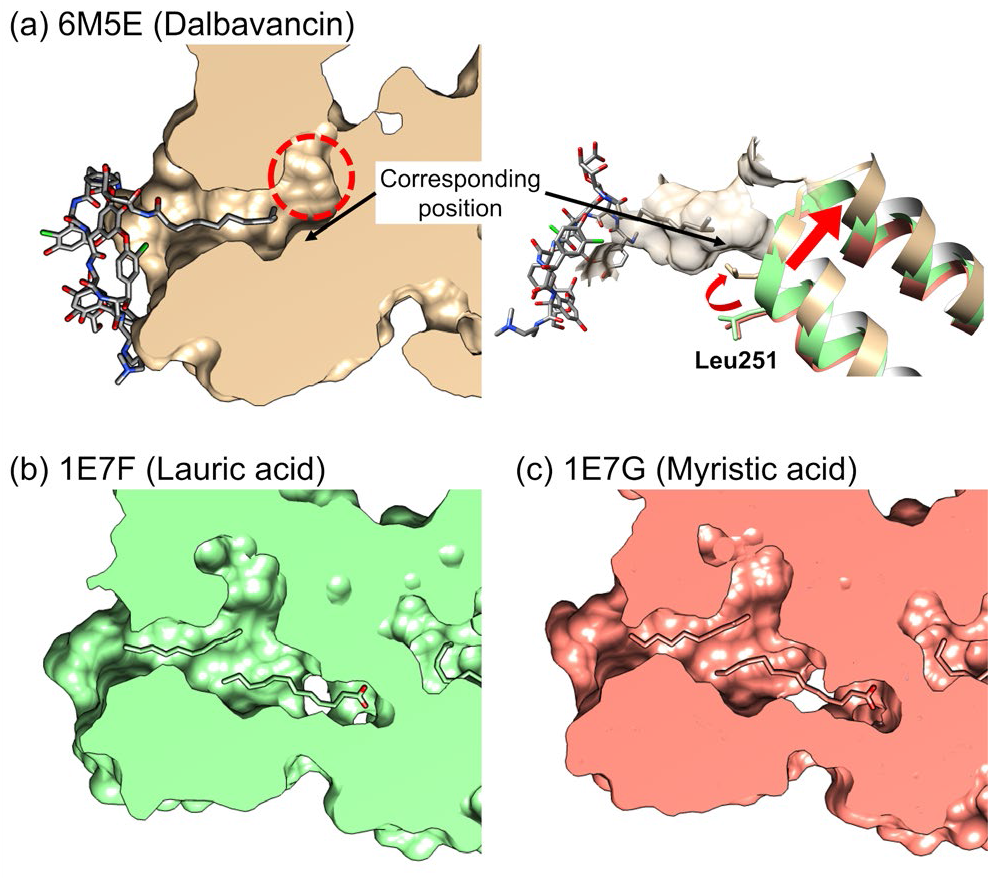
Comparison of the binding pocket in subdomain IA. (a) The surface representation of the binding pocket of dalbavancin and structural alignment of helix 4 of subdomain IIA. (b)The binding pocket of the lauric acids. (c) The binding pocket of the myristic acids.

The cyclic moiety of dalbavancin formed hydrogen bonds with Glu48, Thr52, Asp56, Glu60, and Asn61 (Figure 3b). Importantly, Glu60 was covered by the cyclic moiety, thereby increasing the interaction area between this residue and dalbavancin. Glu60 formed two hydrogen bonds with dalbavancin (Figure 3a and Figure 3b), suggesting that it is a key residue for interaction with the peptide. In this HSA structure, compared to other HSA structures on PDB, Glu60 as well as the surrounding loop region (Ala55 - Ser65) were severely disordered, indicating that this region has high intrinsic flexibility. Nonetheless, clear electron density around Glu60 in the HSA-dalbavancin complex was observed, and this loop region enclosed dalbavancin to increase the interaction area (Figure S2), implicating that dalbavancin may bind to the subdomain IA by an induced-fit mechanism.^28^ In addition, interaction analysis based on the MD trajectory also showed stable hydrogen bonds with Thr52 and Glu60 (Figure 3c and Figure S7). These residues formed hydrogen bonds with nitrogen atoms from dalbavancin backbone in more than 50% of total simulation time. Glu48 formed water-bridged hydrogen bonds with oxygen atoms from dalbavancin backbone or hydroxyl groups more than 30% of the time. The backbone oxygen atom from Lys51 formed hydrogen bonds more than 30% of the time. Moreover, Glu45 formed water-bridged hydrogen bonds with the oxygen atom from the hydrocarbon chain amide group more than 20% of the time; Asn61 and Lys73 also formed hydrogen bonds or water-bridged hydrogen bonds with the same amide group. In summary, dalbavancin is recognized using a combination of both hydrophobic and hydrophilic interactions with the subdomain IA of HSA.

### Binding of other cyclic peptides to albumin and optimization strategies

The structure of HSA in complex with dalbavancin and its binding mechanism were revealed in this study. Despite the fact that cyclic peptides are already used for clinical purposes, this is the first report of a structure of HSA in complex with one of these drugs. Importantly, cyclic peptides harboring a hydrocarbon chain may interact with the same binding site as dalbavancin and via a similar binding mechanism. For example, colistin, which also features a hydrocarbon chain (C8:0), may also bind to this hydrophobic pocket. The dalbavancin-binding pocket has room to potentially accommodate longer hydrocarbon chains, compared to the binding site of myristic acid (C14:0) (Figure 5a). Similarly, caspofungin (C16:0), with a longer hydrocarbon chain than that of dalbavancin, may also interact with this pocket. Daptomycin provides another example of the crucial role of the hydrocarbon chain in these compounds. Interestingly, although the only difference between daptomycin and acetyl-daptomycin is the presence or absence of a hydrocarbon chain, the two display drastically different values of plasma protein binding (85±4.5 % and 12±3.4 %, respectively).^29^ The structure of the HSA-dalbavancin complex implies that daptomycin may also bind to HSA in a similar way as dalbavancin. In fact, the molecular model of HSA-daptomycin complex structure generated by docking simulation showed a similar binding mode (Figure S8). The current view is that acetyl-daptomycin, without the hydrocarbon chain, interacts with HSA only via its cyclic structure, which could lead to the observed lower value of plasma protein binding. This suggests that addition of a hydrocarbon chain could be the simplest way to increase plasma protein binding capacity of cyclic peptides. Other approaches comprise the inclusion of polar, acidic, or basic amino acid residues in cyclic peptides or modification of backbone nitrogen atoms; these strategies may enable cyclic peptides to form hydrogen bonds or salt bridges with amino acids located in helix 3 and 4, such as Glu45, Glu48, Thr52, Asp72, and Lys73. As for the cyclic peptides without hydrocarbon chain, it is unclear how they bind to HSA, considering that the absence of such chain can weaken the interactions between the protein and the drug. Structures of HSA in complex with these drugs will be required to understand their binding mechanism.

## CONCLUSION

In this study, we revealed the crystal structure of the HSA-dalbavancin complex. The crystal structure shows two dalbavancin molecules bound to HSA, one to the subdomain IA and the other to the subdomain IIIB. Based on SAXS and ITC analyses, the binding site for dalbavancin in solution is only subdomain IA; dalbavancin interacts with a hydrophobic pocket in the subdomain IA. Phe70 swings to avoid clashing with the hydrocarbon chain of dalbavancin. Interestingly, the crystal structure also revealed that hydrogen bonds and a large interface area in the loop region of HSA increase the surface interaction between subdomain IA and the cyclic moiety of dalbavancin. In summary, hydrophobic interactions from the hydrocarbon chain and hydrophilic interactions from the cyclic moiety stabilize dalbavancin binding. HSA forms an actual pocket specific for the hydrocarbon chain of dalbavancin. These findings will support optimization of cyclic peptide drug discovery, especially in the ADMET field.

## EXPERIMENTAL SECTION

### Preparation of HSA-dalbavancin complex

Fatty acid and globulin-free HSA was purchased from Sigma-Aldrich (Product # A3782). HSA was dissolved in a buffer containing 50 mM potassium phosphate (pH 7.0) and 150 mM NaCl. Aggregates were removed by centrifugation (15,000 g, 15 min, 4 °C). The protein was purified by size-exclusion chromatography (Superdex 200 10/300) to improve monodispersity, and peak fractions were collected. For crystallization, NaCl was removed from the buffer and the protein was concentrated to 100 mg mL^−1^ using a 50,000 Da molecular mass cut-off concentrator (Millipore). The concentrated protein was incubated with a four-fold molar excess of dalbavancin at 4 °C for 12 h. The microcrystals (10-30 μm) of HSA-dalbavancin (Sigma-Aldrich) complex were obtained in 0.1 M Sodium citrate tribasic dehydrate (pH 5.0) containing 30% (v/v) Jeffamine ED-2001 (pH 7.0) in a week. Crystals were picked up using LithoLoops or nylon loops and flash-frozen in liquid nitrogen without additional cryo-protectant.

### Preparation of HSA domain I-II and domain III

Defatted human albumin domain I-II or domain III was purchased from Albumin Bioscience (Catalog # 9905, # 9903). Domain I-II or III was dissolved in a buffer containing 10 mM HEPES (pH 7.5), 150 mM NaCl. Aggregates were removed by centrifugation (20,000 g, 10 min, 4 °C). The protein was purified by size-exclusion chromatography (Superdex 200 10/300) to improve monodispersity, and peak fractions were collected.

### Crystallization of apo structure

HSA was dissolved in 50 mM potassium phosphate (pH 5.25). Aggregates were removed by centrifugation (15,000 g, 15 min, 4 °C). HSA crystals were grown from a solution containing 100 mg mL^−1^ of protein, 5 mM sodium azide, and 38% (v/v) polyethylene glycol 400 by streak seeding. Large crystals (0.5 × 0.5 × 0.5 mm^3^) were obtained within a day. Crystals were picked up using LithoLoops or nylon loops and flash-frozen in liquid nitrogen without additional cryo-protectant.

### X-ray data collection

Data collection was performed on BL32XU^30^ and BL26B2^31^ at SPring-8. For the complex crystals, 151 small-wedge datasets were collected using Zoo system.^32^ The collected images were processed using KAMO^33^ with XDS^34^. Out of all of the datasets, 123 were selected via correlation coefficient-based clustering using normalized structure factors, followed by merging by XSCALE with outlier rejections implemented in KAMO. The highest resolution of the merged data was 2.8 Å. For the apo crystals, diffraction data were collected on BL26B2. Most crystals diffracted poorly due to the high solvent content (76%, calculated by phenix.xtriage). Therefore, about 100 datasets were automatically collected and processed using DeepCentering^35^ and KAMO. The best crystal diffracted up to 2.6 Å resolution. Details of the data collection are given in Table 1. The structure was solved by phenix.phaser using PDB ID 1AO6.^36^ Refinement was performed with phenix program suite^37,38^ and COOT^39^.

### SAXS data collection and analysis

SAXS data collection was performed using FR-X (Rigaku) to determine the state of assembly of the HSA-dalbavancin complex in solution. HSA was dissolved in a buffer containing 50 mM HEPES (pH 7.0), 50 mM NaCl, and 10% (v/v) DMSO (buffer A). HSA was purified by size-exclusion chromatography (Superdex 200 10/300) and peak fractions were collected. The solute concentrations were determined using Nanodrop ND-1000 (Thermo Fisher) spectrophotometer and adjusted to 5 mg mL^−1^. To collect data on HSA-dalbavancin, 400 μM dalbavancin was added to buffer A. Buffer frames were averaged and subtracted from the sample. Data averaging and reduction were performed using SAXSLab (Rigaku). Rg and I(0) calculations for the bead model were performed using PRIMUS.^40^ Distance distribution functions P(r) were calculated with GNOM.^41^ Ten structures were determined using GASBOR^42^ and a final structure was obtained by averaging each structure using DAMAVER^43^. For comparison of “in crystal” and “in solution” structures of the HSA-dalbavancin complex, the Rg of the crystal structure of the HSA-dalbavancin complex was calculated with CRYSOL^44^.

### Isothermal titration calorimetry (ITC) measurement of dalbavancin and HSA

For isothermal titration calorimetry measurement, full-length HSA, domain I-II, or domain III was prepared as described in the subsections “Preparation of HSA-dalbavancin complex”, and “Preparation of HSA domain I-II and domain III” with slight modifications. The binding stoichiometry of dalbavancin and HSA was evaluated in 50 mM HEPES (pH 7.5) containing 20% (v/v) DMSO on a VP-ITC instrument (MicroCal) at 37 °C and a stirring speed rate of 307 rpm. The 400 μM dalbavancin solution was titrated into 25 μM HSA solution. The binding activity of dalbavancin to domain I-II or domain III was evaluated in 50 mM HEPES (pH 7.5) containing 20% (v/v) DMSO on an iTC200 instrument (Cytiva) at 37 °C. 500 μM of dalbavancin solution was titrated into 25 μM domain I-II or 50 μM domain III. The binding activity of lauric acid to HSA in full length, domain I-II, and domain III were also evaluated in 50 mM HEPES (pH 7.5) containing 20% (v/v) DMSO on an iTC200 instrument (Cytiva) at 37 °C. 900 μM of lauric acid solution was titrated into 10 μM HSA in full length, 10 μM domain I-II, or 30 μM domain III. The measured heat flow was recorded as a function of time and each peak was integrated to determine the total heat generated from each injection. The recorded heat was plotted against the molar ratio of the protein in the cell and the plot was fitted to generate a binding curve by applying a one-site model using the software Origin7. The stoichiometry was estimated from an inflection point of the binding curve.

### Competitive ITC

A thermal competitive assay^45^ with lauric acid was carried out using an iTC200 instrument (Cytiva) at 37 °C in a buffer containing 50 mM HEPES pH 7.5 and 20% (v/v) DMSO. HSA (50 μM) was placed in the cell with or without 370 μM lauric acid, while 500 μM dalbavancin was placed in the syringe. The concentration of lauric acid and dalbavancin was much higher than their *K*_D_ value. The titration consisted of a 5 μL injection from the syringe. The enthalpy associated with the single titration of dalbavancin to HSA was compared in the presence or absence of lauric acid.

### System preparation, equilibration, and molecular dynamics simulation

For MD simulation, we prepared two systems: the HSA-dalbavancin complex and the dalbavancin-removed HSA system. For the HSA-dalbavancin complex system, chain C, chain H, and chain I were used. For the dalbavancin-removed HSA system, only chain C was used. The HSA-dalbavancin complex and dalbavancin-removed HSA were subjected to assignment of bond orders, hydrogenation, creation of zero-order bonds to metal and disulfide bonds, generation of suitable ionization state, and H-bond optimization with Maestro using OPLS_2005 force field.^47^ The preprocessed HSA-dalbavancin complex structure and the dalbavancin-removed HSA structure were immersed in the orthorhombic box with a buffer distance of 10 Å using the simple point charge (SPC) water model. The system was neutralized by adding counter ions (12 Na+ and 10 Na+ ions for the HSA-dalbavancin complex system and the dalbavancin-removed HSA system, respectively) and 0.15 M NaCl. The Desmond MD package^48^ was used to construct the systems, which contained 83,646 atoms and 78,500 atoms per system. The OPLS_2005 force field was used to calculate the interactions between all atoms. The “u-series” algorithm developed at D.E. Shaw Research was used for calculating the long-range Coulombic interactions. The cut-off radius was set to 9 Å for the short-range van der Waals and electrostatic interactions. The initial temperature and pressure were set to 300 K and 1.01325 bar, respectively. The sampling interval during the simulation was set to 10 ps. The temperature and pressure were controlled during the simulations using the Nose-Hoover thermostat^49,50^ and the Martyna-Tobias-Klein method^51^ respectively. A time step of 2.0 fs was used for the simulations. The default protocols of the Desmond package were used for minimization and MD equilibration of the systems. Finally, 200 ns atomistic MD simulations under the NPT ensemble were performed four times by changing the initial velocity.

### HSA-dalbavancin interaction analysis

The trajectory frames obtained from the MD simulations were subjected to Simulation Interaction Diagram tool implemented in the Maestro. This tool defines hydrogen bonds, hydrophobic interactions, π-π stacking, and salt bridge interactions and quantitatively measures the HSA-dalbavancin interactions.

## ASSOCIATED CONTENT

### Supporting Information

Supporting Information is available free of charge at https://pubs.acs.org/doi/##.####/#######

Figure S1. Packing effect of dalbavancins bound to the subdomain IIIB.

Figure S2. View of the loop region (Ala55 – Ser65).

Figure S3. Experimental SAXS curves and Guinier plot.

Figure S4. Competitive ITC of dalbavancin in the presence of lauric acid.

Figure S5. Binding characterization of lauric acid by ITC.

Figure S6. Control titrations in ITC experiments.

Figure S7. The two dimensional schematic diagram of interactions between dalbavancin (chain H) and HSA from MD simulation and the histogram of interactions (Run2, Run3 and Run4).

Figure S8. Top scoring molecular model of the HSA-daptomycin complex structure.

Table S1. Conditions of SAXS data collection, data reduction, analysis and interpretation.

Table S2. Thermodynamic parameters of HSA-dalbavancin interaction by ITC.

The crystal structure of apo form of HSA and HSA-dalbavancin complex have been deposited under PDB accession codes 6M5D and 6M5E, respectively. Authors will release the atomic coordinate and experimental data upon article publication.

## AUTHOR INFORMATION

### Authors

Akinobu Senoo - Department of Chemistry and Biotechnology, School of Engineering, The University of Tokyo, Tokyo 113-8656, Japan;

Satoru Nagatoishi - Institute of Medical Science, The University of Tokyo, Minato-ku, Tokyo 108-8639, Japan;

Masahito Ohue - Department of Computer Science, School of Computing, Tokyo Institute of Technology, Kanagawa, 226-8501, Japan;

Masaki Yamamoto - RIKEN SPring-8 Center, Sayo-gun, Hyogo, 679-5148, Japan and Graduate School of Life Science, University of Hyogo, Hyogo, 678-1297, Japan;

Kouhei Tsumoto - Department of Chemistry and Biotechnology, School of Engineering, Institute of Medical Science and Department of Bioengineering, School of Engineering, The University of Tokyo, Tokyo, 113-8656, Japan;

### Author Contributions

Conceptualization: N.W., S.I., A.S.; Sample preparation: S.I., A.S.; Structure determination by crystallography and SAXS: S.I.; Molecular dynamics: N.W., M.O.; Isothermal titration calorimetry: A.S., S.I., S.N., K.T.; Writing - original draft: N.W., S.I., A.S.; Supervision: N.W., S.I.; Writing - review and editing: N.W., S.I., A.S., S.N., K.T., M.O., M.Y.

\# S.I. and A.S. contributed equally to this work.

### Notes

The authors declare no competing financial interest.

## ACKNOWLEDGMENTS

This research was supported by Platform Project for Supporting Drug Discovery and Life Science Research (Basis for Supporting Innovative Drug Discovery and Life Science Research (BINDS)) from AMED under the Grant Numbers JP19am0101001, JP19am0101094 and JP20am0101094, and by the Japanese Society for the Promotion of Science (JSPS), KAKENHI Grant Number 20K19926. We thank beamline staff members at BL32XU of SPring-8 (Hyogo, Japan) and K. Yamashita for technical help with data collection. We also thank T. Matsumoto for technical help with the SAXS experiment. The numerical calculations were carried out on the TSUBAME3.0 supercomputer at Tokyo Institute of Technology. H. Ago, K. Takeshita, N. Sakai, G. Ueno, and K. Yoshida are acknowledged for their careful reading of the manuscript and valuable comments.

## ABBREVIATIONS

HSA: human serum albumin
PDB: protein data bank
SAXS: small angle x-ray scattering
Rg: radius of gyration
MD: molecular dynamics
FA: fatty acid
ITC: isothermal titration calorimetry
ADMET: absorption, distribution, metabolism, excretion, toxicity

