## Supplemental Information for "Structural basis for binding mechanism of human serum albumin complexed with cyclic peptide dalbavancin"

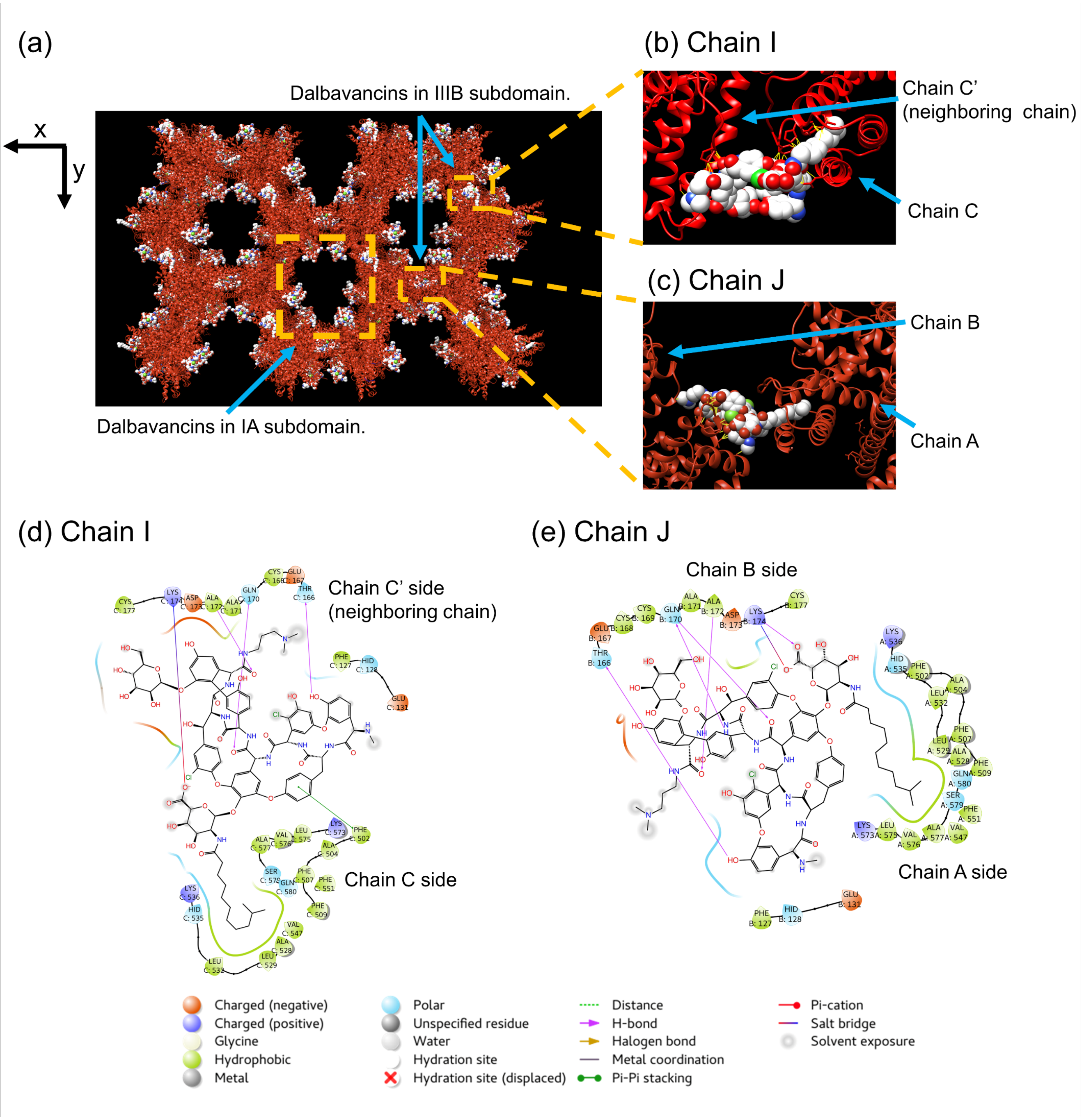


Figure S1. Packing effect of Dalbavancins bound to the subdomain IIIB. (a) Crystal packing of HSA-dalbavancin complex. HSA and Dalbavancins are shown in the ribbon (red) and sphere (white) models, respectively. Although the cyclic moiety of the dalbavancins bound to IA subdomain are located in void area in the crystal, dalbavancins bound to the subdomain IIIB involve crystal packing and tightly interact with neighboring HSA. (b), (c) close-up view of dalbavancin bound to the subdomain IIIB. (d), (e) The two-dimensional schematic diagram of interactions between dalbavancin bound to the subdomain IIIB and HSA. Interactions and amino acid types are represented by their symbols as displayed in the key.

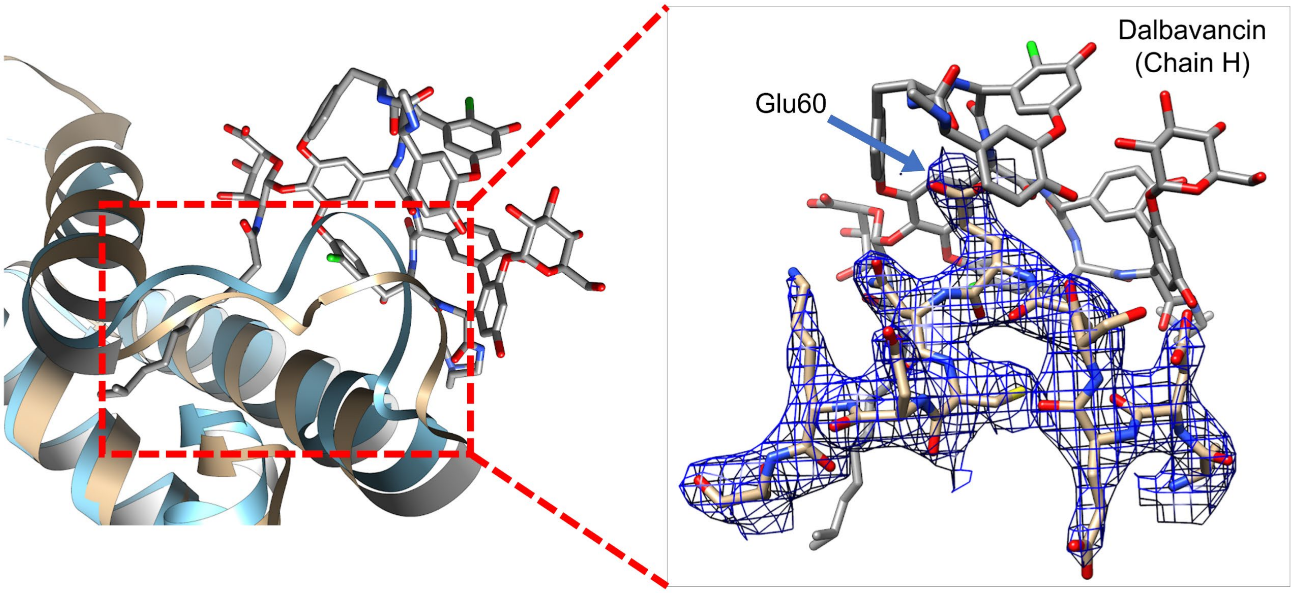


Figure S2. A close-up view of the loop region (Ala55 – Ser65). $2mF_o$-$DF_c$ map of the region contoured at 1.5 σ. Compared to other HSA structures on PDB, electron density of the loop region is clearly observed; Glu60 covers dalbavancin to increase the interaction. This indicates that dalbavancin interacts with HSA via an induced-fit mechanism.

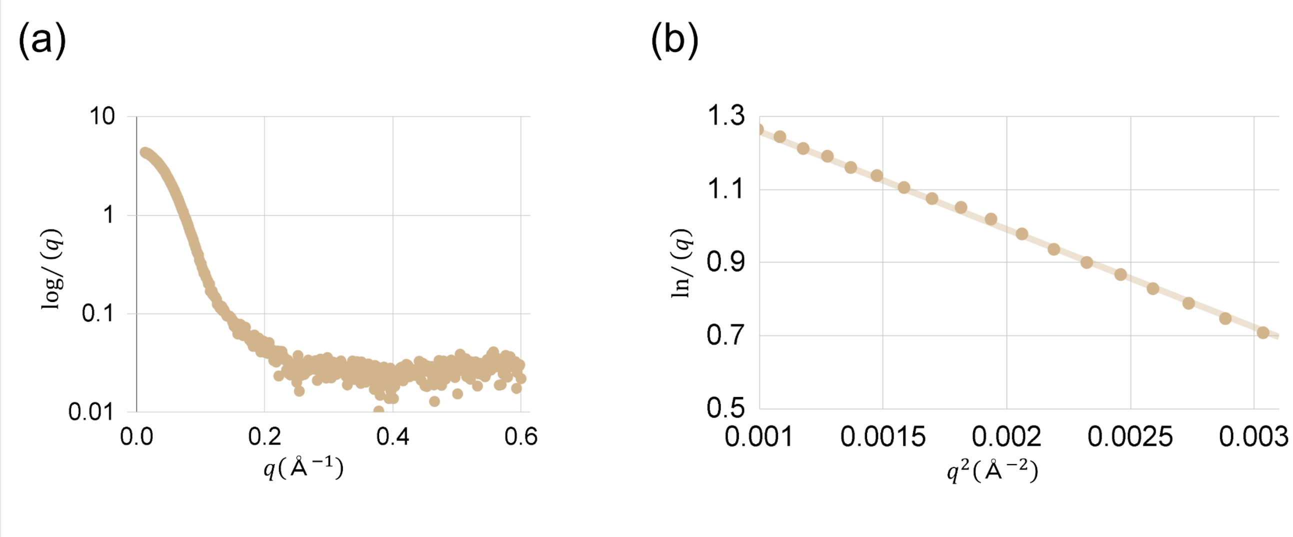


Figure S3. Experimental SAXS curves and Guinier plot of HSA-dalbavancin complex. (a) The experimental scattering curve. (b) The Guinier plot.

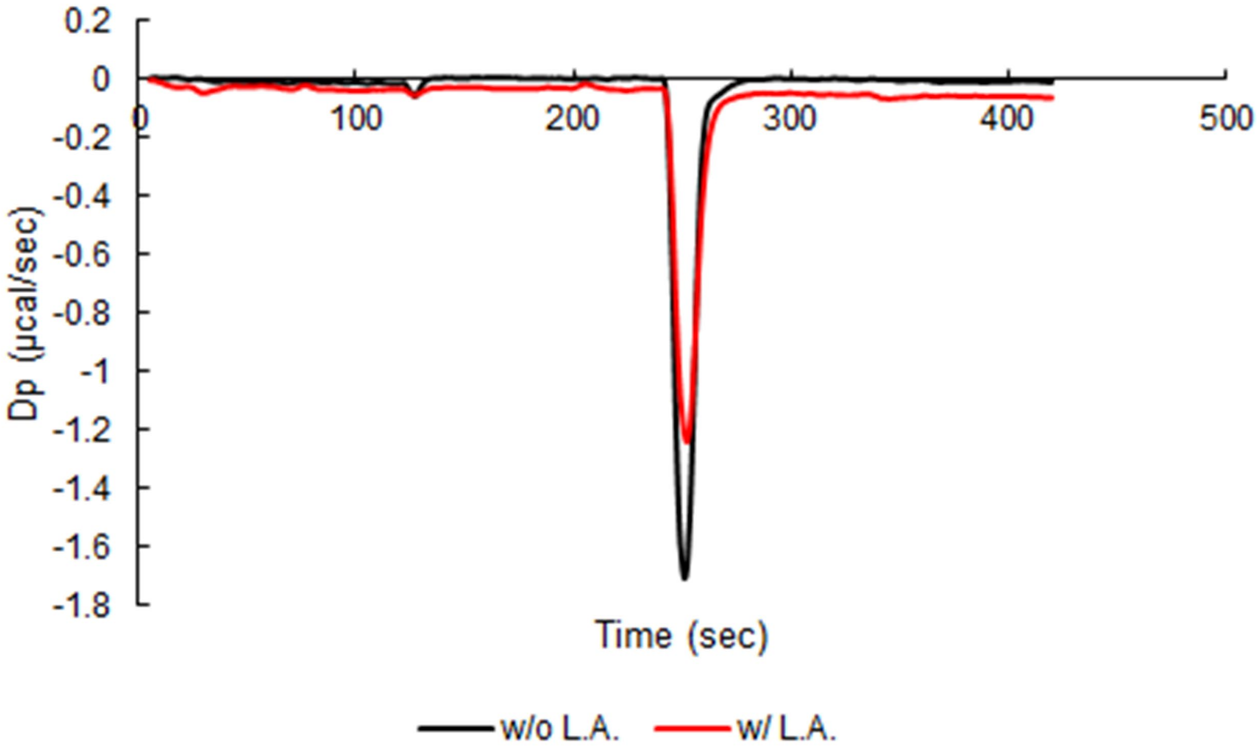


Figure S4. Competitive ITC of dalbavancin in the presence of lauric acid. 500 µM dalbavancin was titrated into the cell that contained 50 µM full-length HSA with or without 350 µM lauric acid (L.A.). Reaction heat upon the single injection is shown in red trace for the condition with lauric acid, and in black trace for the one without lauric acid.

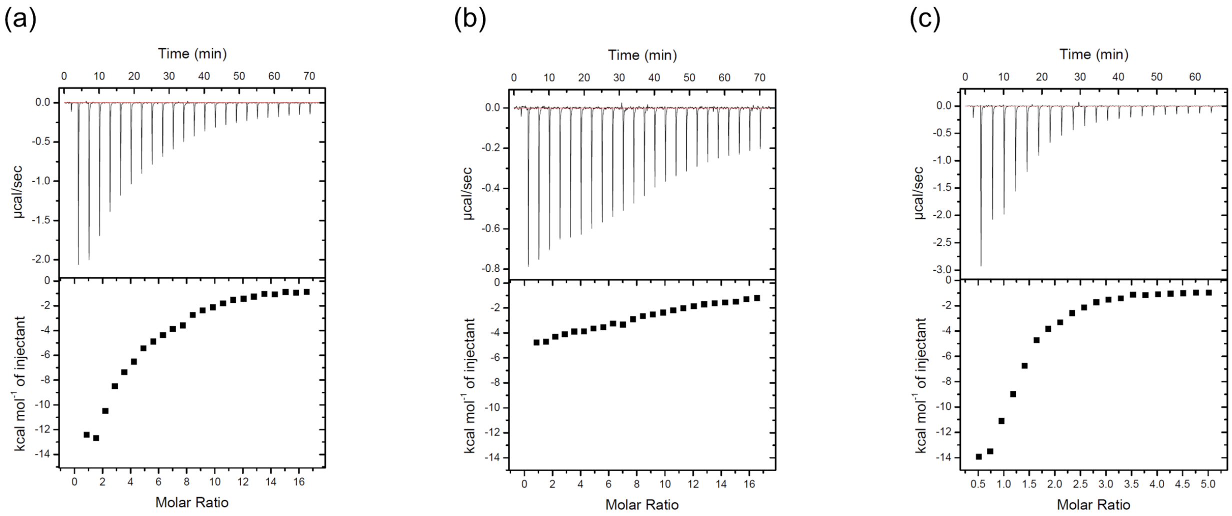


Figure S5. Binding characterization of lauric acid. 900 µM lauric acid was titrated into the cell that contained (a) 10 µM full-length HSA, (b) 10 µM domain I-II, (c) 30 µM domain III. The plot of the first injection is removed in the bottom panel of (c), since the amount of heat upon the injection was higher than that observed in (a), which could have been derived from the unexpected reaction at the artificial interface generated by cleaving the domain III from full-length HSA. The reaction in (a) is roughly divided into two phases, the former phase corresponding to the reaction with domain III and the later phase to the reaction with domain I-II.

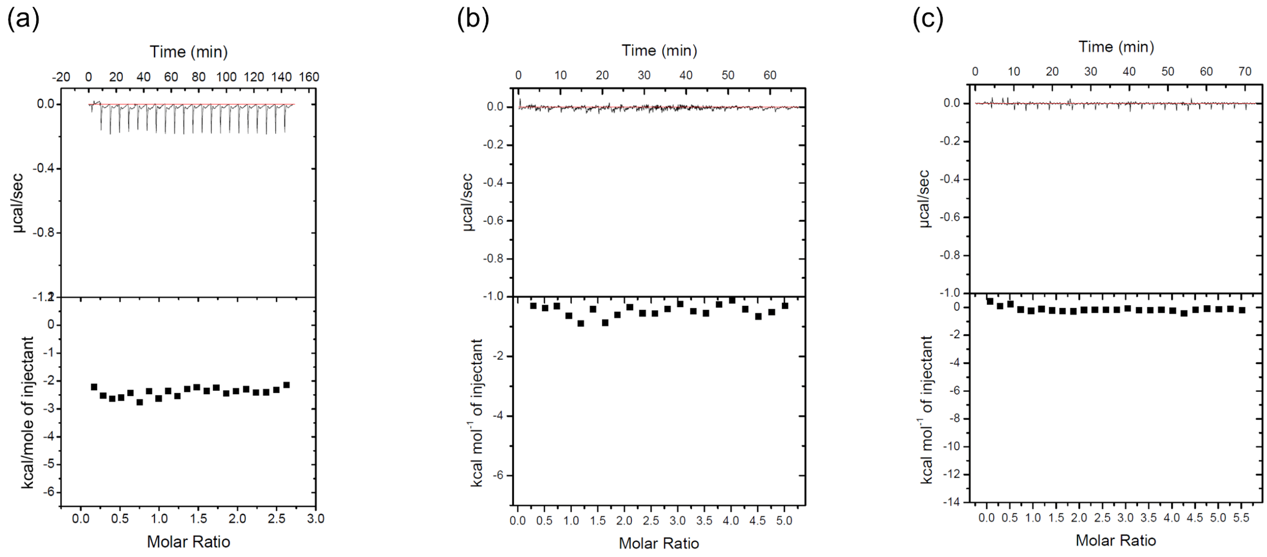


Figure S6. Control titrations of ITC experiments. (a) 400 µM dalbavancin was titrated into the cell that contained only buffer using a VP-ITC instrument. (b) 500 µM dalbavancin was titrated into the cell that contained only buffer using an iTC200 instrument. (c) 900 µM lauric acid was titrated into the cell that contained only buffer using an iTC200 instrument. For all the experiments, 50 mM HEPES (pH 7.5), 20% DMSO buffer was used. These data are not subtracted from Figure 2b-2d or Figure S5, since the dilution heat was small enough and constant through all of the injections.

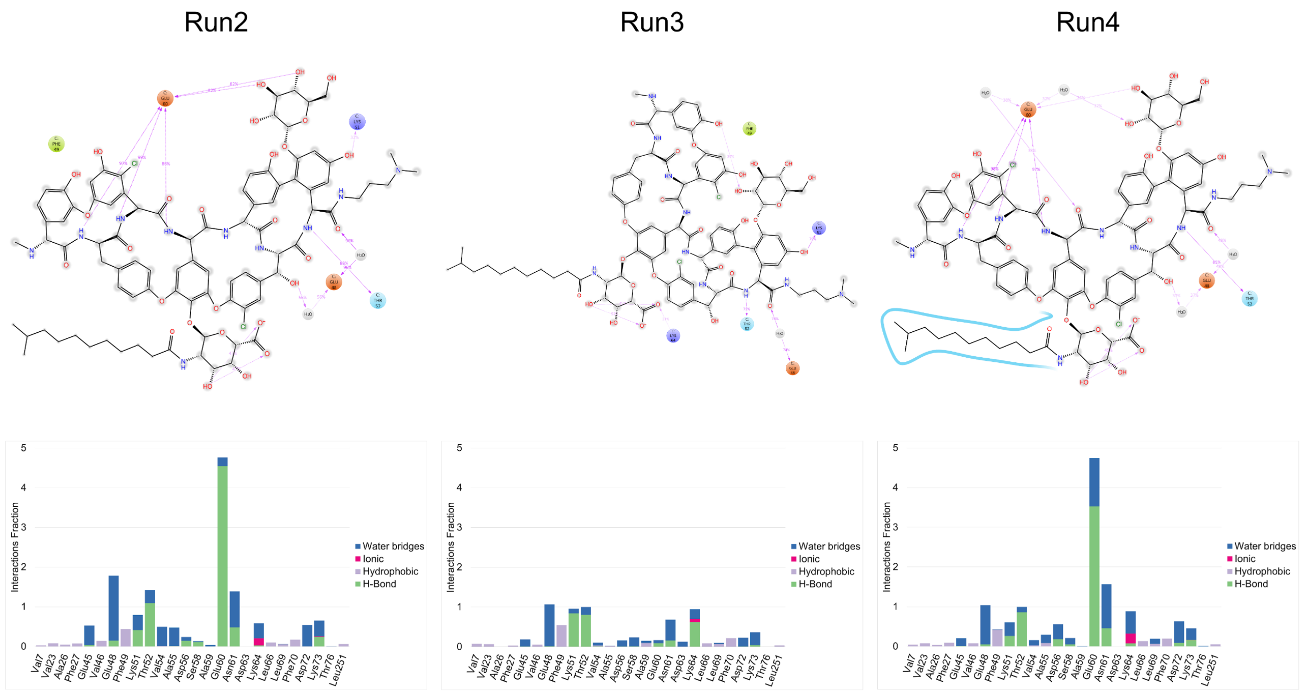


Figure S7. The two-dimensional schematic diagram of the interactions between dalbavancin (chain H) and HSA from MD simulation and the histogram of interactions (Run2, Run3, and Run4). The stacked bar charts are normalized over the course of the trajectory.

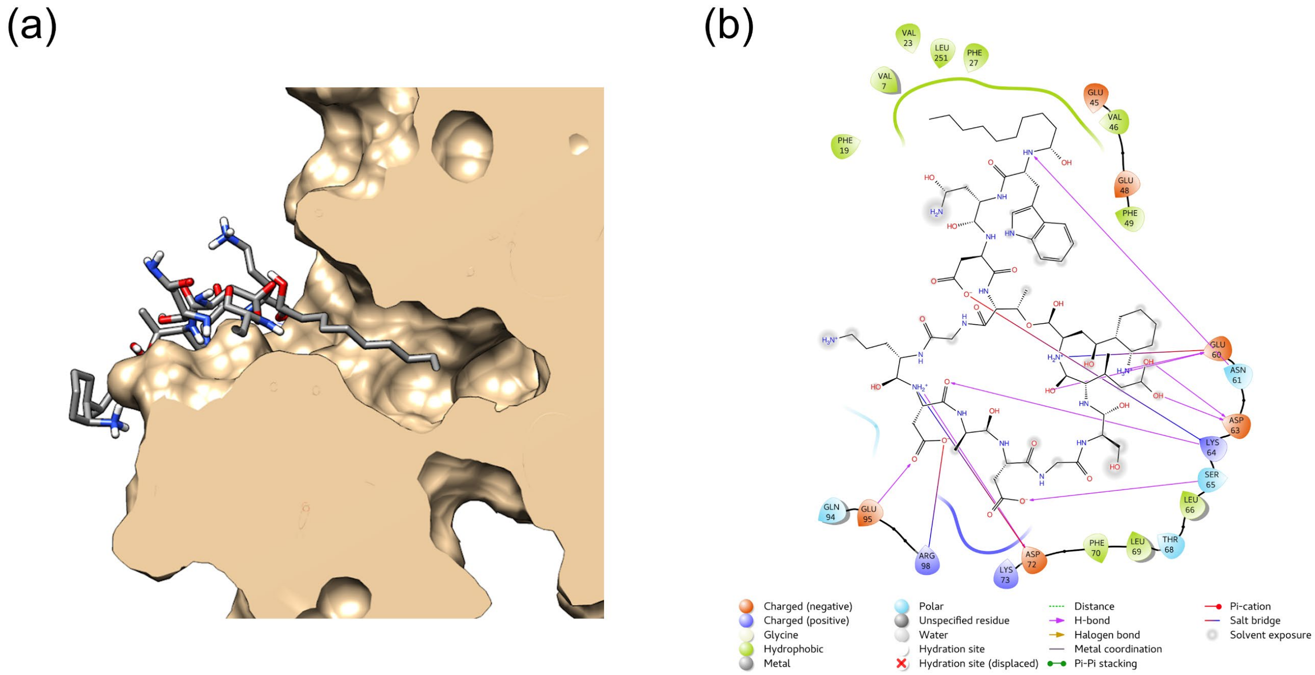


Figure S8. Top scoring molecular model of HSA-daptomycin complex structure. (a) The dalbavancin binding pocket in the IA subdomain (chain C) with daptomycin. (b) The two dimensional schematic diagram of the interactions between the HSA and the daptomycin from docking simulation.

**Table S1. Conditions of SAXS data collection, data reduction, analysis, and interpretation.**

| **SAXS data collection and analysis parameters** | HSA-dalbavancin |
|---|---|
| SAXS system | *BIOSAXS-1000 (Rigaku)* |
| Wavelength (Å) | 1.54178 |
| q range (Å$^{-1}$) | 0.009 - 0.639 |
| Number of frames | 12 |
| Exposure time per frames (minutes) | 15 |
| Sample concentration (mg/mL) | 5 |
| Ligand concentration (uM) | 400 |
| Temperature (°C) | 25 |
| **Structural parameters** | |
| Guinier analysis | |
| I(0) (cm$^{-1}$) | 4.60(1) |
| Rg (Å) | 28.2(1) |
| q range (Å$^{-1}$) | 0.11 - 0.048 |
| qRg max | 1.3 |
| Dmax (Å) | 83.00 |
| Coefficient of correlation, $R^2$ | 0.999 |
| **Shape model-fitting results** | |
| *GOSBOR (default parameters, 10 calculations)* | |
| q range for fitting | 0.013 - 0.600 |
| Symmetry, anisotropy assumptions | P1, none |
| $\chi 2$ range | 1.299 - 1.831 |
| $\chi 2$ average | 1.522 |
| **Software employed for SAXS data reduction, analysis and interpretation** | |
| SAXS data reduction | *PRIMUSqt from ATSAS* |
| Basic analysis: Guinier | *SAXSlab* |
| Shape modelling | *GASBOR* |
| Three-dimensional graphic model representations | *chimera* |

**Table S2. Thermodynamic parameters of HSA-dalbavancin interaction by ITC.**

| | $K_D$ (μM) | $\Delta H$ (kcal $mol^{-1}$) | $-T\Delta S$ (kcal $mol^{-1}$) | $n$ |
|---|---|---|---|---|
| full length | 11.6 | -8.9 | 1.9 | 0.84 |
| domain I-II | 5.0 | -7.7 | 0.17 | 1.27 |

Modeling of the HSA-daptomycin complex using docking simulation

Protein preparation, ligand preparation and docking simulation

Our HSA-dalbavancin complex structure (PDB ID: 6M5E) was used as a docking target. This structure was subjected to protein preparation processes as described in the main text. Docking simulations were conducted for the dalbavancin binding site, only for the subdomain IA, of the prepared HSA-dalbavancin complex structure in the absence of dalbavancin. A grid box with dimensions of $20 \times 20 \times 20$ Å$^3$ was generated to maintain the dalbavancin binding site. To prepare the ligand structures, the NMR solution state structures of daptomycin (PDB ID: 1T5M[1]) were subjected to Ligprep[2] to generate tautomeric and ionization states, ring conformations and stereoisomers. In total, 7680 structures were generated. Docking simulation experiments were carried out using Glide[3,4] standard-precision mode and 7680 daptomycin structures were docked into the dalbavancin binding site of HSA. The docking poses were evaluated based on the Glide docking score.

Model of HSA-daptomycin complex structure

Docking simulations generated 4310 docking poses and the top three poses had docking scores of -11.30, -10.38 and -10.37. These models of HSA-daptomycin complex structure showed similar binding mode to HSA-dalbavancin complex structure (Figure S8a). Hydrocarbon chain of daptomycin interacted with non-polar amino acids including Val7, Phe19, Val23, Phe27, Val46, Phe49, Leu66, Leu69, Phe70 and Leu251; the Cyclic moiety also interacted with polar residues located in the loop region (Ala55 – Ser65). Especially, Glu60 formed hydrogen bonds with the backbone of cyclic moiety and a salt bridge with the side chain of Kyn13 (non-proteinogenic amino acid in daptomycin), Asp63 formed hydrogen bonds with the side chain of threo-Glu12, Lys64 formed a salt bridge with side chain of Asp3 and Ser65 formed a hydrogen bond with side chain of Asp9 (Figure S8b).
